# HCLC-FC: a novel statistical method for phenome-wide association studies

**DOI:** 10.1101/2022.03.14.484203

**Authors:** Xiaoyu Liang, Xuewei Cao, Qiuying Sha, Shuanglin Zhang

**Author notes:** **Corresponding author** (SZ).

## Abstract

The emergence of genetic data coupled to longitudinal electronic medical records (EMRs) offers the possibility of phenome-wide association studies (PheWAS). In PheWAS, the whole phenome can be divided into numerous phenotypic categories according to the genetic architecture across phenotypes. Currently, statistical analyses for PheWAS are mainly univariate analyses, which test the association between one genetic variant and one phenotype at a time. In this article, we derived a novel and powerful multivariate method for PheWAS. The proposed method involves three steps. In the first step, we apply the bottom-up hierarchical clustering method to partition a large number of phenotypes into disjoint clusters within each phenotypic category. In the second step, the clustering linear combination method is used to combine test statistics within each category based on the phenotypic clusters and obtain p-values from each phenotypic category. In the third step, we propose a new false discovery rate (FDR) control approach. We perform extensive simulation studies to compare the performance of our method with that of other existing methods. The results show that our proposed method controls FDR very well and outperforms other methods we compared with. We also apply the proposed approach to a set of EMR-based phenotypes across more than 300,000 samples from UK Biobank. We find that the proposed approach not only can well-control FDR at a nominal level but also successfully identify 1,244 significant SNPs that are reported to be associated with some phenotypes in the GWAS catalog. Our open-access tools and instructions on how to implement HCLC-FC are available at https://github.com/XiaoyuLiang/HCLCFC.

**Author summary:** As a complementary approach to genome-wide association studies, phenome-wide association studies (PheWAS) have been an efficient tool for testing associations between genetic variations and a wide range of phenotypes utilizing all available phenotypic information. For instance, the first PheWAS has demonstrated that rs3135388 on *HLA-DRB1* associated with atrial fibrillation and multiple sclerosis. A challenging step in performing large-scale multiple testing of PheWAS is to control the false discovery rate (FDR). In this work, we propose a novel and powerful multivariate method, HCLC-FC, to test the association between a genetic variant with a large number of phenotypes simultaneously controlling FDR. Within each phenotypic category, a newly proposed method clusters phenotypes into different groups and the combined test statistic within each category based on the phenotypic clusters has an asymptotic distribution which avoids the computational burden of simulation. Furthermore, the newly developed FDR controlling process is based on p-values and does not depend on test statistics. Therefore, it is more general and can be applied to other multiple testing procedures to control FDR.

## Introduction

Genome-wide association studies (GWAS) have emerged as a common and powerful tool for investigating the genetic architecture of human disease over the last ten years [1, 2]. In general, the conventional GWAS focus on a single phenotype or disease, aiming to identify the association between single nucleotide polymorphisms (SNPs) and a univariate phenotype or disease [2]. Over the last decade, numerous disease- and trait-associated common SNPs have been successfully identified by using statistical methods of GWAS. However, these common SNPs explain only a small portion of inheritable phenotypic variance for human disease [3]. Therefore, recent genetic studies have suggested that many genetic loci appear to harbor variants associated with multiple phenotypes [4].

To date, many software packages, such as PLINK, Gen/ProbABEL, MaCH, SNPTEST, and FaST-LMM, have been developed to support GWAS [5-11]. However, GWAS suffer from important shortcomings. First of all, GWAS usually focus on a pre-defined and limited phenotypic domain and ignore the potential power gained through the use of intermediate phenotypes that may more closely reflect a gene’s mechanism, as well as the association between genetic variation and multiple phenotypes [12, 13]. Moreover, it is difficult to reach the threshold of statistical significance by GWAS due to the burden of multiple comparisons conducted; only those associations with a p-value less than 5×10^−8^ are considered statistically significant. GWAS have difficulty in explaining a significant portion of the predicted phenotypic heritability even though a significant number of SNPs are identified [14]. Lastly, the genotype-phenotype association is assessed for millions of SNPs one by one, false-positive results may easily arise due to large-scale multiple testing. Therefore, a large sample size is needed to achieve the optimal statistical power and minimize the spurious associations. Furthermore, the replication of the significant loci in independent populations is necessary according to the GWAS criteria [15].

Recently, large-scale DNA databanks linked to longitudinal electronic medical records (EMRs) offer the possibility of phenome-wide association studies (PheWAS) and have been proposed as an approach for rapidly generating large, diverse cohorts for discovery and replication of genotype-phenotype associations [16-18]. In most EMR systems, the whole phenome can be classified into numerous phenotypic categories according to genotypic and phenotypic information such as phenotype similarity [17], genetic architecture [19], and disease network [20]. As a complementary approach to GWAS, PheWAS investigate the association between SNPs and a diverse range of phenotypes. By utilizing all available phenotypic information and all genetic variants in the estimation of associations between genotype and phenotype, a broader picture of the relationship between genetic variation and networks of phenotypes is possible [19]. In summary, GWAS use a phenotype-to-genotype strategy, beginning with a specific phenotype or disease; PheWAS reverse this paradigm by using a genotype-to-phenotype approach, starting with a genotype to test for associations over a wide spectrum of human phenotypes [14].

We are motivated primarily by PheWAS, which aim to assess associations between SNPs and a diverse range of phenotypes. Many of the issues that arise in this setting also occur elsewhere, for example, in clinical trials, the outcomes of cardiovascular risk may include hospitalization, stroke, heart failure, myocardial infarction, cardiac arrest, disability, and death [21]. Therefore, the statistical framework and results given here have a potential for wider application. In 2019, Sha et al. [22] developed the Clustering Linear Combination (CLC) method that combines univariate test statistics for jointly analyzing multiple phenotypes in association analysis. CLC has been theoretically proved to be the most powerful test among all tests with certain quadratic forms if the phenotypes are clustered correctly. It is not only robust to different signs of means of individual statistics but also reduces the degrees of freedom of the test statistics. Therefore, the CLC method can be applied to PheWAS. However, due to the unknown number of clusters for a given data, the final test statistic of the CLC method is the minimum p-value among all p-values of the test statistics obtained from each possible number of clusters [23] and a simulation procedure is used to estimate the p-value of the final test statistic which would be time-consuming especially in the PheWAS setting.

In this article, we derived a novel and powerful multivariate method, which we referred to as HCLC-FC (Hierarchical Clustering Linear Combination with False discovery rate Control), to test the association between a genetic variant with a large number of phenotypes. The HCLC-FC method is applicable to EMR-based PheWAS. In PheWAS, the whole phenome can be classified into numerous phenotypic categories according to genotypic and phenotypic information, and each category contains a certain number of phenotypes. The proposed method (HCLC-FC) involves three steps. In the first step, we use the bottom-up Hierarchical Clustering Method (HCM) [24] to partition a large number of phenotypes into disjoint clusters within each category. In the second step, we apply the CLC method to combine test statistics within each phenotypic category based on the phenotypic clusters and obtain p-values from each phenotypic category. In the third step, we develop a false discovery rate (FDR) control approach based on a large-scale association testing procedure with theoretical guarantees for FDR control under flexible correlation structures [12]. Using extensive simulation studies, we evaluate the performance of the proposed method and compare the power of the proposed method with the powers of three commonly used methods in association studies using multiple phenotypes. These three methods include Multivariate Analysis of Variance (MANOVA) [25], joint model of Multiple Phenotypes (MultiPhen) [26], and Trait-based Association Test that uses Extended Simes procedure (TATES) [27]. Our simulation studies show that the proposed method outperforms the other three methods for different within-group and between-group phenotypic correlation structures we consider. Furthermore, the existing methods using our proposed FDR control procedure can control FDR efficiently. We also evaluate the performance of HCLC-FC through a set of 1,869 EMR-based phenotypes based on the International Classification of Diseases, 10^th^ Revision (ICD-10 code, Data-Field 41202), across more than 300,000 samples from the UK Biobank, where these phenotypes can be classified into 260 ICD-10 level 1 blocks. The real data analysis results show that HCLC-FC can well control the type I error rate and can identify 1,244 SNPs that have previously been reported in the GWAS catalog.

## Materials and methods

### Statistical methods

Consider a sample with *n* unrelated individuals for a PheWAS, indexed by *i* = 1,2,…,*n*. Each individual has the phenome with *K* phenotypes. The *K* phenotypes can be divided into *M* phenotypic categories, indexed by *m* = 1,…,*M*. Suppose that there are *K*_*m*_ phenotypes in the *m*^*th*^ category, where *m* = 1,2,…,*M* and 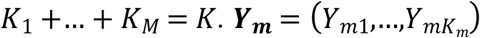 is the *n*× *K*_*m*_ phenotype matrix of *m*^*th*^ phenotypic category, where *Y*_*mk*_ = (*y*_1*mk*_,…,*y*_*nmk*_)^*T*^ and *y*_*imk*_ is the *k*^*th*^ phenotype in the *m*^*th*^ category of the *i*^*th*^ individual. Denote *X* = (*x*_1_,…,*x*_*n*_)^*T*^ as the genotypic score of *n* individuals at a genetic variant of interest, where *x*_*i*_ ∈ {0, 1, 2} is the number of minor alleles that the *i*^*th*^ individual carries at a genetic variant. We are interested in simultaneously testing the collection of *M* hypotheses *H*_0*m*_: the *m*^*th*^ phenotypic category is not associated with the genetic variant of interest.

We assume that there are no covariates. If there are covariates, such as, gender, age, BMI, and top principal components to adjust for population stratification, we adjust both phenotype and genotype values for the covariates using the method applied by Price et al. [28] and Sha et al. [29]. That is, if there are *p* covariates, *z*_*i*1_,…,*z*_*ip*_, for the *i*^*th*^ individual, we adjust both phenotype and genotype values for the covariates through linear models

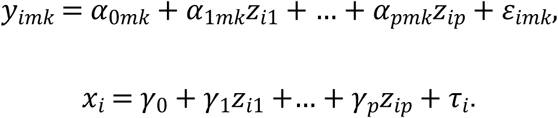

In this article, we derived a novel and powerful multivariate method for PheWAS, which is referred to as HCLC-FC. The proposed method (HCLC-FC) involves three steps. In the first step, we use the bottom-up HCM [24] to partition *K*_*m*_ phenotypes into *L*_*m*_ disjoint clusters within each category, where *m* = 1,…,*M*. In the second step, we apply the CLC [22] to combine test statistics within each category. The CLC test statistic with *L*_*m*_ clusters follows a chi-square distribution with *L*_*m*_ degrees of freedom. We then obtain the p-value of the CLC test statistic for each phenotypic category. In the third step, we propose an FDR control approach based on the method proposed by Cai et al. [12]. FDR is widely used to claim significance for high-dimensional correlated data. However, most of the existing methods of FDR cannot accurately estimate FDR due to different directions of genetic effects on different phenotypes. Recently, Cai et al. (Cai, et al., 2019) developed a method to evaluate FDR that works well for single-phenotype PheWAS. However, Cai’s method is based on test statistics which are difficult to extend to test statistics for multiple phenotypes. Instead of using test statistics, we propose a new approach to evaluate FDR which is based on p-values and does not depend on test statistics. In the following sections, we give a detailed approach for each step.

#### Step 1: HCM to partition phenotypes in each phenotype category

For the *m*^*th*^phenotypic category, we partition *K*_*m*_ phenotypes into *L*_*m*_ disjoint clusters. Denote ***D***_***m***_ = **1** − **Σ**_***m***_ with entries 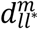 as the dissimilarity matrix, where **Σ**_***m***_ is *K*_*m*_ × *K*_*m*_ correlation matrix of ***Y***_***m***_ for the *m*^*th*^ phenotypic category and 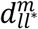 is the dissimilarity value between *l*^*th*^ and *l*^\**th*^ phenotypes. The HCM is based on the agglomerative clustering algorithm. In agglomerative clustering, all the phenotypes are a cluster of their own, and we merged pairs of clusters until they form a single cluster. In each iteration, we merge two clusters that have the smallest value of the average dissimilarity 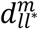 between all phenotypes in two clusters and define the smallest average dissimilarity *h*_*b*_ as the height of the *b*^*th*^ iteration. HCM uses the established principle in Bühlmann et al. [30] to determine the number of clusters for each phenotypic category. That is, the number of clusters *L*_*m*_ is identified at the 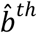 iteration, where 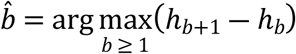

#### Step 2: CLC to test the association between phenotypes in each category and a genetic variant

For each phenotypic category, we apply the CLC method [22] to combine test statistics among the *L*_*m*_ clusters. We use *T*_*mk*_ to denote the score test statistic to test the null hypothesis *H*_0*mk*_:*β*_1*mk*_ = 0 (the *k*^*th*^ phenotype in the *m*^*th*^ phenotypic category is not associated with the genetic variant) under the generalized linear model *y*_*imk*_ = *β*_0*mk*_ + *β*_1*mk*_*x*_*i*_ + *ε*_*imk*_, where *k* = 1,…,*K*_*m*_. So *T*_*mk*_ is given by 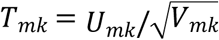, where 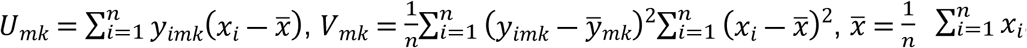, and 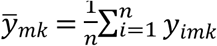. If we let 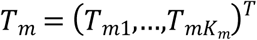 be the test statistic vector that contains score test statistics for each phenotype in the *m*^*th*^ phenotypic category and let ***B***_***m***_ be a *K*_*m*_× *L*_*m*_ matrix with the indicator entry *b*_*kl*_ = 1 if the *k*^*th*^ phenotype belongs to the *l*^*th*^ cluster and *b*_*kl*_ = 0 otherwise. Then the CLC test statistic for the *L*_*m*_ clusters in the *m*^*th*^ phenotypic category is given by 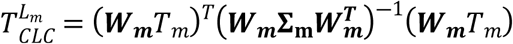, where 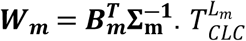 follows a chi-square distribution with *L* _*m*_ degrees of freedom. We denote *p*_*m*_ as the p-value of 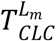.

#### Step 3: Threshold for FDR-controlling

The method proposed by Cai et al. [12] is based on test statistics which are hard to extend to other test statistics. Therefore, in this step, we develop a new approach to evaluate FDR which is based on p-values. In the second step, the p-value for the test statistic in the *m*^*th*^ category for *m* = 1,…,*M* can be obtained. In this step, we propose a new multiple testing FDR controlling procedure by thresholding the p-values {*p*_*m*_:*m* = 1,…,*M*}. Under the null hypothesis, each *p*_*m*_ follows a uniform distribution *U*(0,1). Let *t*, 0 ≤ *t* ≤ 1, be a rejection threshold so that *H*_0*m*_ is rejected if and only if *p*_*m*_ ≤ *t*. For any given threshold *t*, 0 ≤ *t* ≤ 1, the false discovery proportion (FDP) based on a random sample is given by

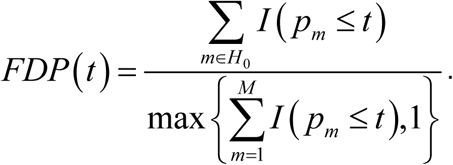

To maximize the power of the test or equivalently the rejection rate among ℋ_1_ while maintaining an FDP level of *α*, the optimal threshold *t* is 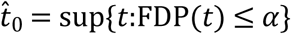. The key to empirically controlling the FDP is to find a good estimate of the numerator ∑_*m*_∈*H*_0_ *I*(*p*_*m*_ ≤ *t*). Using the idea in Cai et al. [12], we estimate the numerator by ∑_*m*_∈*H*_0_ *I*(*p*_*m*_ ≤ *t*) ≈ *m*_0_*G*(*t*), where *m*_0_ is the number of categories under the null hypothesis and we can use *M* to estimate *m*_0_ due to the sparsity in the number of alternative hypotheses in many real data applications, and *G*(*t*) = *P*(*U*(0,1) ≤ *t*) = *t*.

For a given nominal FDR level *α* ∈ (0,1), we reject *H*_0*i*_ whenever 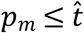, where

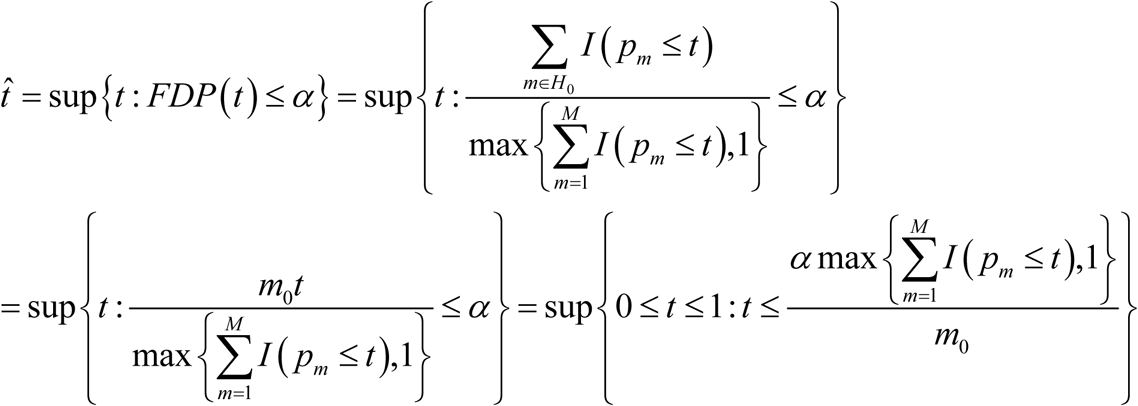

### Comparison of methods

We compare the performance of the proposed method HCLC-FC with those of MultiPhen [26], MANOVA [25], and TATES [27]. To evaluate the FDR-controlling performance, MANOVA, MultiPhen, and TATES are first applied to each category. Then, we apply the third step of HCLC-FC to the three methods to control FDR, which are referred to as MANOVA-FC, MultiPhen-FC, and TATES-FC. That is, we not only compare the performance of different methods for joint analysis of multiple phenotypes but also compare the performance of different methods with the newly developed FDR-controlling process.

In the following sections, we will estimate the FDR and power of each method. FDR is estimated by FDP, 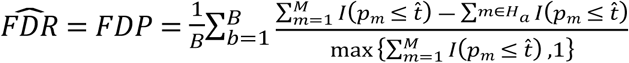, where *B* is the number of replications, *H*_*a*_ is the alternative hypothesis, *p*_*m*_ is the p-value of the test statistic for the *m*^*th*^ phenotypic category, *m* = 1,…,*M*, and 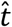 is the threshold estimated by HCLC-FC in step 3. The power of each method is the probability of correctly rejecting *H*_0_; it is estimated by 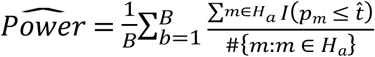.

### Simulation study

To evaluate the FDRs and powers of the proposed method, we generate genotypes according to the minor allele frequency (MAF) of a genetic variant and assume Hardy Weinberg equilibrium. Then, we generate *K* phenotypes by the following models similar to the models used by Sha et al. [2019] and Liang et al. [2018] [22, 24]. Suppose there are *M* categories and there are 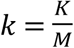 phenotypes in each category. Let *y*_*m*_ = (*y*_*m*1_,…,*y*_*mk*_)^*T*^ denote the phenotypes in the *m*^*th*^ category. We assume

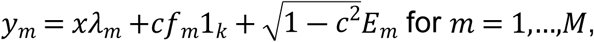

where *x* is the genotype score at the variant of interest; *λ*_*m*_ = (*λ*_*m*1_,…,*λ*_*mk*_)^*T*^ is the vector of effect sizes of the genetic variant on phenotypes in the *m*^*th*^ category; *f* = (*f*_1_,…,*f*_*M*_)^*T*^ ∼*MVN*_*M*_(0,**Σ**_***f***_), **Σ**_***f***_ = (1 − *ρ*_*f*_)***I*** + *ρ*_*f*_***A***, *ρ*_*f*_ is a constant to define the phenotypic correlation between phenotypic categories, ***A*** is an *M* × *M* matrix with elements of 1, and ***I*** is an *M* × *M* identity matrix; *c* is a constant; *E*_1_,…,*E*_*M*_ are independent and *E*_*m*_∼*MVN*_*k*_(0,**Σ**_***e***_) with **Σ**_***e***_ = (*σ*_*ij*_), where 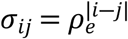 and *ρ*_*e*_ is constant to define the phenotypic correlation within each phenotypic category.

Based on equation (1), we consider the following six models. In these six models, the correlation between the *i*^*th*^ and *j*^*th*^ phenotypes within each category is 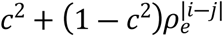, and between categories is *c*^2^*ρ*_*f*_. We set *M* =100 for Model 1-3 and *M* = 50 for Model 4-6.

**Model 1**: There are *M* =100 categories and genotypes impact on only one category. Let *λ*_1_ = … = *λ*_*M*−1_ = 0 and *λ*_*M*_ = *β*(1,…,*k*)^*T*^.

**Model 2**: There are *M* =100 categories and genotypes impact on two categories.

Let 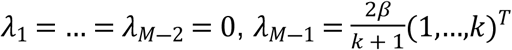, and 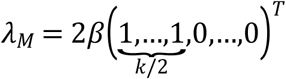.

**Model 3**: There are *M* =100 categories and genotypes impact on three categories. Let 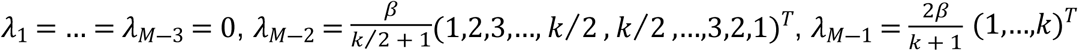 and 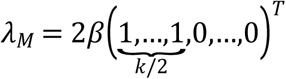.

**Model 4**: Same as Model 1, but there are *M* = 50 categories.

**Model 5**: Same as Model 2, but there are *M* = 50 categories.

**Model 6**: Same as Model 3, but there are *M* = 50 categories.

## Results

### Simulation results

In our simulation studies, we estimate the p-values of all test statistics using their asymptotic distributions. We first set *ρ*_*f*_ = 0.2, *ρ*_*e*_ = 0.3, *c*^2^ = 0.5, and *K* = 1,000, 2,000 for comparing the performance of different methods for joint analysis of multiple phenotypes, in other words, we consider the proposed FDR-controlling method and compare the performance of HCLC-FC, MANOVA-FC, MultiPhen-FC, and TATES-FC. For FDR evaluation, we consider different numbers of phenotypes, different sample sizes, different values of effect size, and different models.

The estimated FDRs of the four methods are summarized in **Table 1** and **Table 2**. From these tables, we can see that all methods using our FDR control procedure control their respective targeted error rates very well, which indicates applying our new FDR-controlling procedure on the entire collection of hypotheses can control the rate of FD of associated genetic variants as well as the expected value of the average proportion of FD of phenotypic categories influenced by such variants.

**Table 1.** The estimated FDR of the four tests under the six models for 1,000 phenotypes (*K* = 1,000). MAF is 0.3. The sample size (*n*) is 2,000. *ρ*_*f*_ = 0.2, *ρ*_*e*_ = 0.3, and *c*^2^ = 0.5. *β* is the effect size. FDR is evaluated using 200 replicated samples at a nominal FDR level of 5%. All estimated FDR are within the 95% confidence interval (0.0198, 0.0802).

| Model | $\beta$ | Method | | | |
| --- | --- | --- | --- | --- | --- |
|  |  | HCLC-FC | MANOVA-FC | MultiPhen-FC | TATES-FC |
| 1 | 0.012 | 0.038 | 0.039 | 0.048 | 0.041 |
|  | 0.014 | 0.045 | 0.029 | 0.033 | 0.037 |
|  | 0.016 | 0.039 | 0.049 | 0.047 | 0.048 |
| 2 | 0.050 | 0.034 | 0.048 | 0.047 | 0.044 |
|  | 0.060 | 0.041 | 0.042 | 0.041 | 0.037 |
|  | 0.070 | 0.049 | 0.053 | 0.045 | 0.073 |
| 3 | 0.050 | 0.049 | 0.037 | 0.048 | 0.063 |
|  | 0.090 | 0.047 | 0.043 | 0.046 | 0.048 |
|  | 0.130 | 0.048 | 0.057 | 0.057 | 0.063 |
| 4 | 0.005 | 0.043 | 0.063 | 0.063 | 0.035 |
|  | 0.006 | 0.047 | 0.061 | 0.049 | 0.045 |
|  | 0.007 | 0.041 | 0.050 | 0.049 | 0.065 |
| 5 | 0.050 | 0.048 | 0.052 | 0.056 | 0.030 |
|  | 0.060 | 0.048 | 0.047 | 0.044 | 0.034 |
|  | 0.070 | 0.042 | 0.038 | 0.048 | 0.050 |
| 6 | 0.050 | 0.035 | 0.064 | 0.065 | 0.040 |
|  | 0.090 | 0.055 | 0.039 | 0.049 | 0.034 |
|  | 0.130 | 0.047 | 0.044 | 0.043 | 0.046 |

**Table 2.**
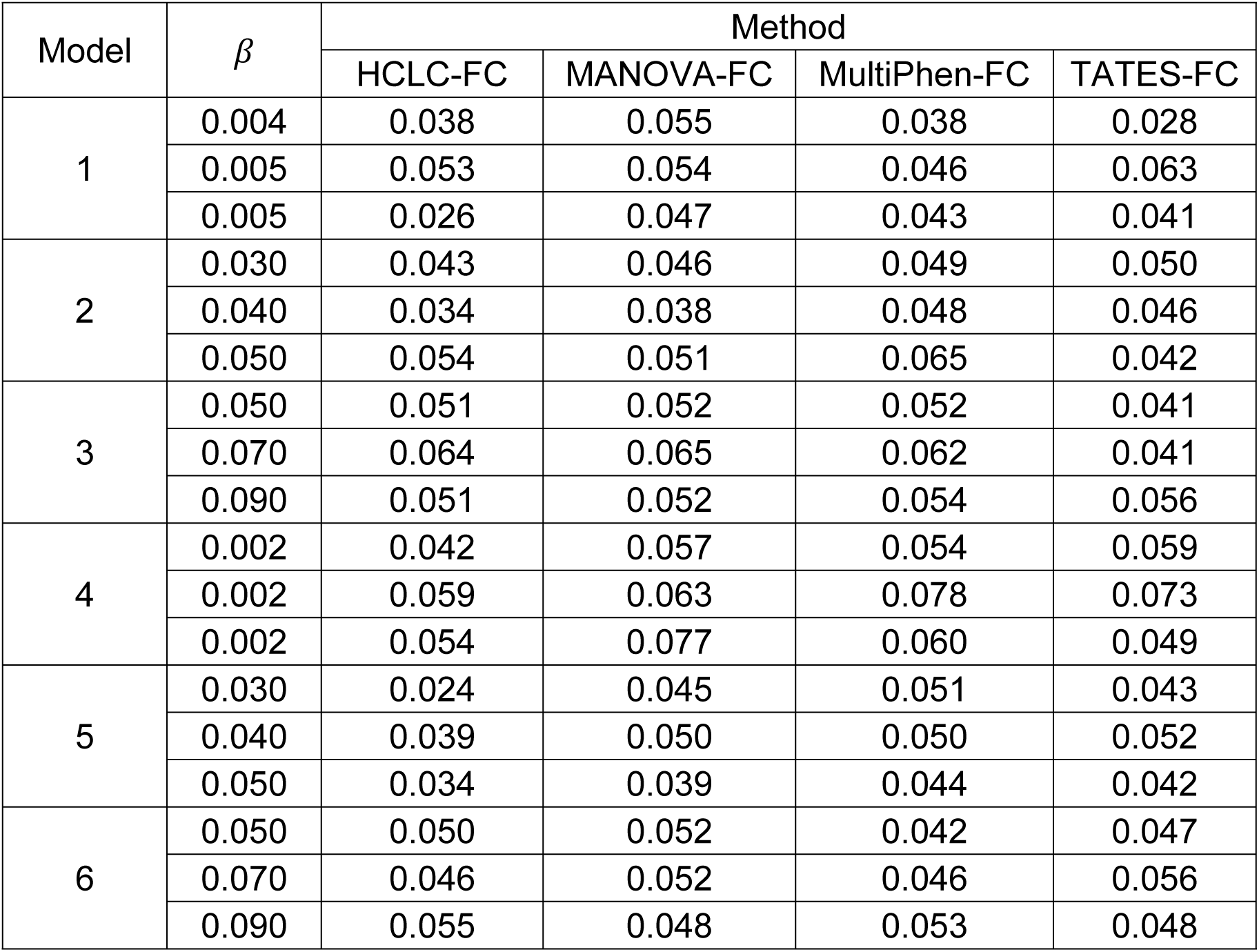
The estimated FDR of the four tests under the six models for 2,000 phenotypes (*K* = 2,000). MAF is 0.3. The sample size (*n*) is 4,000. *ρ*_*f*_ = 0.2, *ρ*_*e*_ = 0.3, and *c*^2^ = 0.5. *β* is the effect size. FDR is evaluated using 200 replicated samples at a nominal FDR level of 5%. All estimated FDR are within the 95% confidence interval (0.0198, 0.0802).

| Model | $\beta$ | Method | | | |
| --- | --- | --- | --- | --- | --- |
|  |  | HCLC-FC | MANOVA-FC | MultiPhen-FC | TATES-FC |
| 1 | 0.004 | 0.038 | 0.055 | 0.038 | 0.028 |
|  | 0.005 | 0.053 | 0.054 | 0.046 | 0.063 |
|  | 0.005 | 0.026 | 0.047 | 0.043 | 0.041 |
| 2 | 0.030 | 0.043 | 0.046 | 0.049 | 0.050 |
|  | 0.040 | 0.034 | 0.038 | 0.048 | 0.046 |
|  | 0.050 | 0.054 | 0.051 | 0.065 | 0.042 |
| 3 | 0.050 | 0.051 | 0.052 | 0.052 | 0.041 |
|  | 0.070 | 0.064 | 0.065 | 0.062 | 0.041 |
|  | 0.090 | 0.051 | 0.052 | 0.054 | 0.056 |
| 4 | 0.002 | 0.042 | 0.057 | 0.054 | 0.059 |
|  | 0.002 | 0.059 | 0.063 | 0.078 | 0.073 |
|  | 0.002 | 0.054 | 0.077 | 0.060 | 0.049 |
| 5 | 0.030 | 0.024 | 0.045 | 0.051 | 0.043 |
|  | 0.040 | 0.039 | 0.050 | 0.050 | 0.052 |
|  | 0.050 | 0.034 | 0.039 | 0.044 | 0.042 |
| 6 | 0.050 | 0.050 | 0.052 | 0.042 | 0.047 |
|  | 0.070 | 0.046 | 0.052 | 0.046 | 0.056 |
|  | 0.090 | 0.055 | 0.048 | 0.053 | 0.048 |

To compare the power of HCLC-FC with that of MANOVA-FC, MultiPhen-FC, and TATES-FC, we consider different numbers of phenotypes, different sample sizes, different models, and different genetic effect sizes. The power of the four tests at an FDR level of 5% for 1,000 phenotypes and 2,000 phenotypes are shown in **Fig 1** and **Fig 2**, respectively. According to the power comparison results, we summarize the following conclusions. (1) HCLC-FC outperforms MANOVA-FC, MultiPhen-FC, and TATES-FC consistently for all models we consider; HCLC-FC is the most powerful test no matter the effect sizes show no groups (Model 1 and 4) or show some groups (Model 2, 3, 5, and 6) within the categories impacted by the SNP; (2) MANOVA-FC and MultiPhen-FC have similar power and are more powerful than TATES-FC for all models we consider.

**Fig 1.**
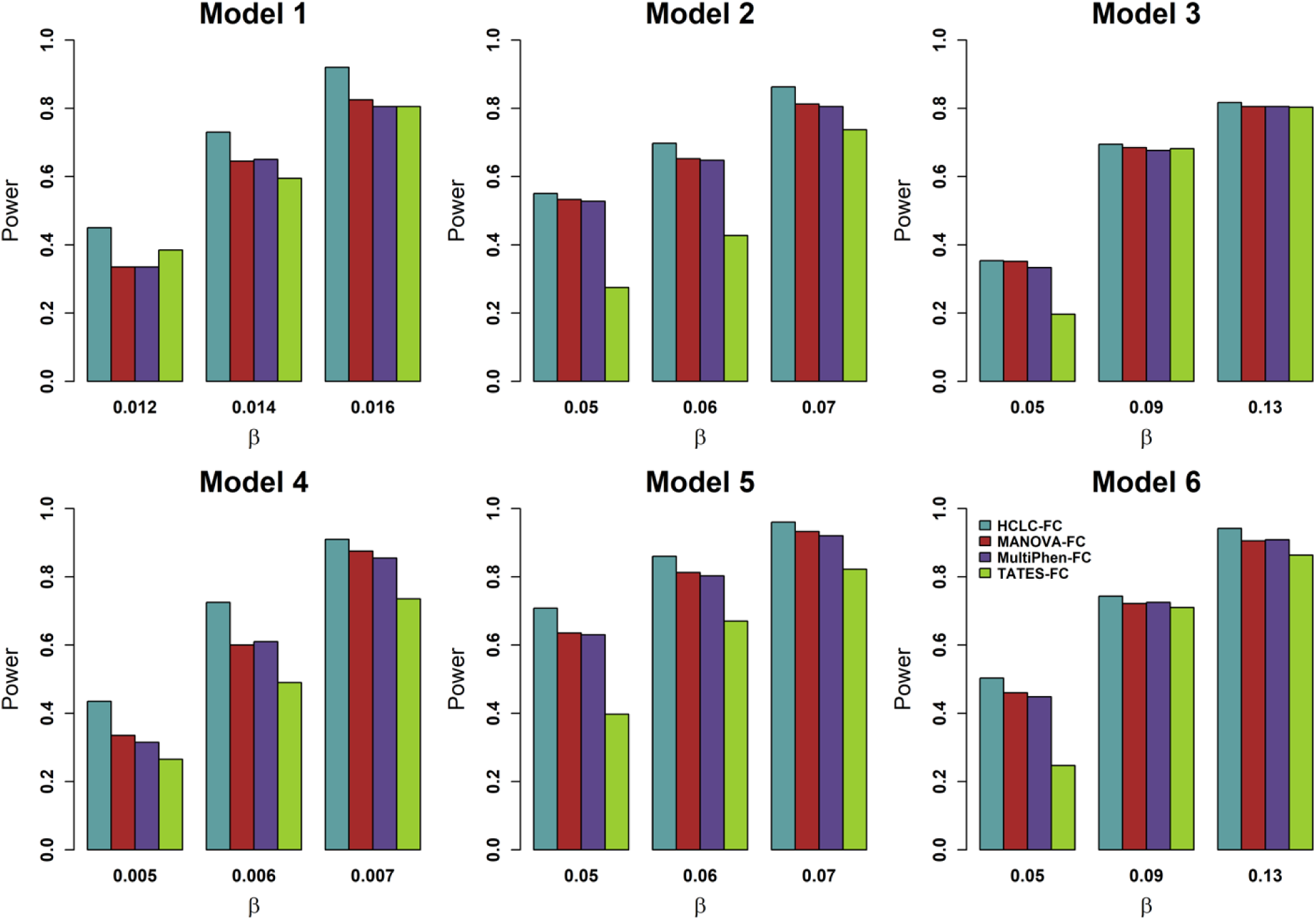
Power comparisons of the four tests for the power as a function of effect size (*β*) under the six models for 1,000 phenotypes (*K* = 1,000). MAF is 0.3. The sample size (*n*) is 2,000. *ρ*_*f*_ = 0.2, *ρ*_*e*_ = 0.3, and *c*^2^ = 0.5. The power of all of the four tests is evaluated using 200 replicated samples at a nominal FDR level of 5%.

**Fig 2.**
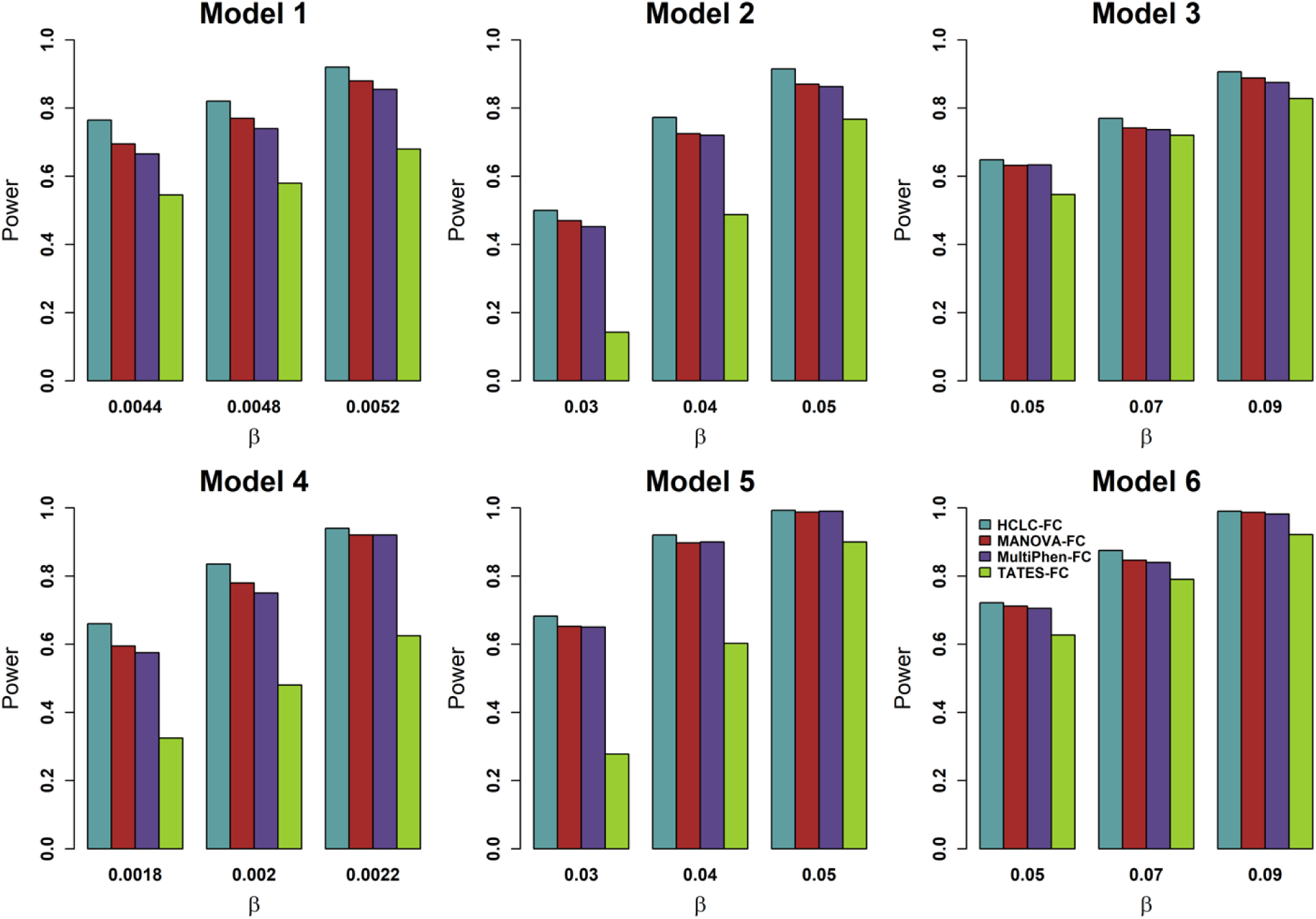
Power comparisons of the four tests for the power as a function of effect size (*β*) under the six models for 2,000 phenotypes (*K* = 2,000). MAF is 0.3. The sample size (*n*) is 4,000. *ρ*_*f*_ = 0.2, *ρ*_*e*_ = 0.3, and *c*^2^ = 0.5. The power of all of the four tests is evaluated using 200 replicated samples at a nominal FDR level of 5%.

In addition to considering power as a function of genetic effect size, we further evaluate power as a function of the correlation between phenotypic categories *ρ*_*f*_ (**S1 Fig**), correlation within each phenotypic category *ρ*_*e*_ (**S2 Fig**), and the constant *c*^2^ in the model (**S3 Fig**) and fix genetic effect size and the other two parameters each time.

Based on **S1**-**S3 Figs**, we can conclude that HCLC-FC outperforms MANOVA-FC, MultiPhen-FC, and TATES-FC consistently for different correlations between phenotypic categories and correlations within each phenotypic category. For power as a function of *c*^2^ (**S3 Fig**), HCLC-FC is either the most powerful test (Model 1, 2, 4, 5, and 6) or comparable with the most powerful test (In Model 3, *c*^2^ = 0.3). The powers of HCLC-FC, MANOVA-FC, and MultiPhen-FC slightly decrease with the increasing correlation between phenotypic categories and correlation within each phenotypic category (**S1 and S2 Figs**). The powers of HCLC-FC MANOVA-FC, and MultiPhen-FC increase with increasing the constant *c*^2^ in the models, but the power of TATES-FC decreases as increasing the constant *c*^2^ (**S3 Fig**).

In summary, the proposed HCLC-FC method is more powerful than the other three methods we compare with under the six models for different within-category and between-category phenotypic correlation structures. The existing methods using our proposed FDR control procedure can control FDR at a given nominal level.

### Real data applications

The UK Biobank is a population-based cohort study with a wide variety of genetic and phenotypic information [31]. It includes ∼ 500K people from all around the United Kingdom who were aged between 40 and 69 when recruited in 2006-2010 [32]. Genotype and phenotype data from the UK Biobank have 488,377 participants with 784,256 variants on chromosomes 1-22 [33]. The preprocess of genotype is achieved by quality control (QC) which is performed on both genotypic variants and samples using PLINK 1.9 [34] (https://www.cog-genomics.org/plink/1.9/). We summarize the QC procedures in **S4 Fig**. In QC, we filter out genetic variants with variant-based missing rates larger than 5%, p-values of Hardy-Weinberg equilibrium exact test less than 10^−6^, and MAF less than 5%. We also filter out individuals with sample-based genotype missing rates larger than 5% and individuals without sex. After QC, there are 250,850 SNPs and 466,501 individuals remaining in the following analysis.

In this study, we define phenotypes using ICD-10 codes, a standardized coding system for defining disease status as well as for billing purposes [35]. After truncating each full ICD-10 code to the UK Biobank ICD-10 level 2 code (https://biobank.ndph.ox.ac.uk/showcase/field.cgi?id=41202), we generate a total of 1,869 unique phenotypes with the names of these phenotypes being the unique truncated ICD codes. For each individual, we denote the EMR-based phenotype for that individual as “1” if a corresponding truncated ICD code ever appears, otherwise, we denote the EMR-based phenotype as “0”. To ensure the individuals in our analysis are from the same ancestry, we first restrict individuals to be the individuals who self-report themselves from a white British ancestry and have very similar ancestry based on a principal component (PC) analysis of genotypes [36]. To avoid the low quality of phenotype data, we exclude individuals who are marked as outliers for heterozygosity or missing rates and have been identified to have ten or more third-degree relatives or closer. Finally, we also exclude individuals that are recommended for removal by the UK Biobank. After preprocessing the phenotype data, there are 337,285 individuals left (details described in **S4 Fig**). It is worth noting that some individuals violate multiple criteria, therefore, the total number of individuals we start with minus the number of individuals that need to be removed does not necessarily equal the number of individuals we keep.

There are 260 blocks based on the UK Biobank ICD-10 level 1 code, therefore, 1,869 phenotypes from the UK Biobank ICD-10 level 2 can be classified into 260 blocks (*M* = 260). We further limit SNPs of interest to those SNPs reaching the genome-wide significance threshold 5×10^−8^. On Oct. 21^st^, 2019, the GWAS catalog (https://www.ebi.ac.uk/gwas/) contains a total of 90,428 data entries covering 3,153 publications of 61,613 SNPs which contains 29,297 significant SNPs. Among 250,850 SNPs obtained from the UK Biobank after QC, there are 3,267 SNPs matched with those significant SNPs in GWAS Catalog. After preprocessing procedures, individuals with both genotype and phenotype information are used in our study. There is a total of 322,607 individuals across 3,267 common SNPs and 1,869 case-control phenotypes which are classified into 260 blocks. Furthermore, we adjust each phenotype by thirteen covariates, including age, sex, genotyping array, and the first 10 PCs [37].

Based on the results shown in **Table 1** and **Table 2**, we know that HCLC-FC, MultiPhen-FC, MANOVA-FC, and TATES-FC can control targeted FDR under all of the simulation models. However, in the UK Biobank data, most of the phenotypes have extremely unbalanced case-control ratios, where the case-control ratios of 1,869 phenotypes are ranged 3.10×10^−6^ to 1.87×10^−1^. Meanwhile, many widely used approaches for joint analysis of multiple phenotypes produce inflated type I error rates for such extremely unbalanced case-control phenotypes [38]. Notably, our proposed FDR control method assumes that the p-value of the test statistic in the *m*^*th*^ category,*p*_*m*_, for *m* = 1,…,*M*, follows a uniform distribution *U*(0,1). Therefore, we first evaluate the distributions of the p-values under the null hypothesis for each of the four methods based on the UK Biobank data by permutation procedures. For each of the four tests, we randomly permute genotypes for each of the 3,267 SNPs. After permutation, 3,267 SNPs have no association with each of the 260 phenotypic blocks. Therefore, we consider 260 blocks and 3,267 SNPs as 260× 3,267 = 849,420 replicated samples. For each replicated sample, we apply four tests for testing the association between each permuted SNP and each phenotypic block. **S5 Fig** shows the histogram of p-values and QQ plot for uniform distribution for each method based on 849,420 replicated samples. In the histogram of p-values, the red dashed line represents the theoretical frequency (849,420/25 ≈ 33,977) for the standard uniform distribution. The frequencies of the p-values of the HCLC method are the only ones that approach the theoretical frequency. Apparently, the frequencies of MultiPhen’s p-values appear to be higher than the expected frequency from uniform distribution when they are closed to 1, and the p-values from 0 to 0.7 have similar frequencies that are smaller than the expected frequency from a uniform distribution. The frequencies of the p-values of MANOVA are higher than the expected frequency when the p-values are near 0 or 1. Contrarily, those of TATES are higher when the p-values belong to (0.35,0.75] or near 0. We also calculate the genomic inflation factor (*λ*) and show the observed and expected p-values from the standard uniform distribution in quantile-quantile (QQ) plots for each method.

In general, the genomic inflation factor *λ* should be close to 1 if the p-values fall within the standard uniform distribution [39]. In the QQ plots of **S5 Fig**, our proposed HCLC method forms a line that’s roughly straight and *λ* = 0.99, indicating that the p-values based on 849,420 replicated samples come from the standard uniform distribution. In contrast, *λ* = 0.58 for MultiPhen and *λ* = 0.84 for MANOVA, where the sample quantiles of these methods deviate from the theoretical quantile. Even though the genomic inflation factor of TATES is equal to 0.97 which is pretty satisfactory, the sample quantiles fluctuate around the theoretical quantiles lightly which is not as good as our proposed HCLC method.

Since the other three methods do not satisfy the uniform distribution assumption of the p-values under the null hypothesis, we only apply HCLC-FC to test the association between each of the 3,267 SNPs and each of the 260 phenotypic blocks. By controlling the FDR at the 5% level, HCLC-FC identifies 1,244 out of 3,267 SNPs that are significantly associated with at least one phenotypic block. **Table 3** shows the top nine SNPs that are significantly associated with multiple phenotypic blocks identified by the HCLC-FC method. We use SNP rs3129716 as an example. rs3129716 is mapped to gene *HLA-DQB1* and *MTCO3P1*. By controlling the FDR at 5%, the FDR threshold is 5.96×10^−3^. Using this threshold, HCLC-FC identifies 28 phenotypic blocks significantly associated with this SNP. Based on the GWAS catalog, 16 out of 28 phenotypic blocks (bold-faced) are reported to be significantly associated with this SNP.

**Table 3.** The top nine SNPs that are associated with multiple phenotypic blocks identified by the HCLC-FC method based on the UK Biobank data. The information of the phenotypic blocks can be found in https://biobank.ndph.ox.ac.uk/showcase/field.cgi?id=41202. The bold-faced blocks indicate the associations with the corresponding SNP reported in the GWAS catalog. The number under the rs-number of SNP represents the total number of phenotypic blocks identified. FDR threshold is calculated at a nominal FDR level of 5%.

| SNPs | Mapped Gene(s) | FDR Threshold | Phenotypic Blocks |
| --- | --- | --- | --- |
| rs3117582<br>(29) | *APOM* | 5.96E-03 | **C15-C26**, C81-C96, D50-D53, D60-D64, E10-E14, E15-E16, E20-E35, E70-E90, G50-G59, H00-H06, H30-H36, I70-I79, J40-J47, K20-K31, K40-K46, K50-K52, K70-K77, M15-M19, M20-M25, N00-N08, N20-N23, N30-N39, N40-N51, Q00-Q07, R00-R09, R30-R39, R50-R69, Y90-Y98, Z40-Z54 |
| rs3129716<br>(28) | *HLA-DQB1*,<br>*MTCO3P1* | 5.96E-03 | **C15-C26**, **C81-C96**, **D50-D53**, D60-D64, **D65-D69**, E15-E16, **E20-E35**, **E70-E90**, **G50-G59**, H00-H06, **H25-H28**, H30-H36, |
|  |  |  | **H55-H59**, I70-I79, **J40-J47**, K20-K31, K40-K46, **K50-K52**, **L10-L14**, **L40-L45**, M15-M19, **M30-M36**, **N00-N08**, N40-N51, **R00-R09**, R10-R19, R30-R39, **R50-R69** |
| rs389884<br>(27) | *STK19* | 5.58E-03 | **C15-C26**, C81-C96, D50-D53, D60-D64, E10-E14, E15-E16, E20-E35, E70-E90, G50-G59, H00-H06, **H30-H36**, I70-I79, **J40-J47**, K40-K46, K50-K52, K70-K77, M15-M19, M20-M25, **N00-N08**, N20-N23, N30-N39, N40-N51, R00-R09, R30-R39, R50-R69, Y90-Y98, **Z40-Z54** |
| rs3134942<br>(25) | *NOTCH4* | 5.19E-03 | C81-C96, **D50-D53**, D60-D64, **E10-E14**, E15-E16, **E20-E35**, **E70-E90**, G50-G59, H00-H06, **H30-H36**, I70-I79, **J40-J47**, K40-K46, **K50-K52**, L20-L30, M15-M19, **M20-M25**, M45-M49, M50-M54, M80-M85, N00-N08, N30-N39, R00-R09, R30-R39, R50-R69 |
| rs3130288<br>(23) | *ATF6B* | 4.81E-03 | C81-C96, D50-D53, D60-D64, E10-E14, E15-E16, E20-E35, G50-G59, H00-H06, H30-H36, I70-I79, **J40-J47**, K40-K46, K50-K52, M15-M19, **M20-M25**, N00-N08, N20-N23, N30-N39, N40-N51, R00-R09, R30-R39, R50-R69, Y90-Y98 |
| rs3094005<br>(22) | *MICB* | 4.62E-03 | **C15-C26**, **C81-C96**, D50-D53, D60-D64, D80-D89, E10-E14, E15-E16, **E20-E35**, G50-G59, H00-H06, H30-H36, **I70-I79**, **J40-J47**, K40-K46, **K50-K52**, K70-K77, **M20-M25**, N00-N08, N20-N23, R00-R09, R30-R39, **Z40-Z54** |
| rs9270493<br>(22) | *HLA-DRB1*,<br>*HLA-DQA1* | 4.62E-03 | **C69-C72**, D50-D53, D60-D64, **D65-D69**, D80-D89, **E10-E14**, E15-E16, **E20-E35**, G50-G59, **H00-H06**, H30-H36, **I70-I79**, K20-K31, **K50-K52**, L00-L08, **L40-L45**, **M05-M14**, **M20-M25**, **N00-N08**, N80-N98, R10-R19, R30-R39 |
| rs35242582<br>(22) | *HLA-DQA1* | 4.62E-03 | A75-A79, **C15-C26**, **C43-C44**, **C60-C63**, **C73-C75**, **C76-C80**, D00-D09, **D50-D53**, E00-E07, E10-E14, **E20-E35**, **G35-G37**, **K50-K52**, L55-L59, **L80-L99**, **M05-M14**, M45-M49, M50-M54, N40-N51, **N80-N98**, O20-O29, U00-U49 |
| rs1480380<br>(22) | *HLA-DMB*,<br>*HLA-DMA* | 4.42E-03 | D10-D36, D50-D53, D60-D64, E00-E07, E10-E14, E15-E16, **E70-E90**, G10-G14, G50-G59, H00-H06, H30-H36, K20-K31, K40-K46, K50-K52, L10-L14, M15-M19, **M20-M25**, **N00-N08**, N80-N98, R00-R09, V30-V39, X60-X84 |

To visualize the associations identified by our proposed HCLC-FC method, we use two sets of phenotypic blocks, the diseases of the circulatory system (I00-I99) and the malignant neoplasms (C00-C97). **Fig 3** shows the associations between SNPs (red circle) and the diseases of the circulatory system phenotypic blocks (I00-I99; blue square) identified by the HCLC-FC method. There are nine phenotypic blocks within the set of the diseases of the circulatory system (I00-I99), such as acute rheumatic fever (I00-I02) and chronic rheumatic heart diseases (I05-I09). We can see from **Fig 3** that many SNPs are associated with one phenotypic block, however, some SNPs are associated with multiple phenotypic blocks. For example, 108 SNPs are associated with the phenotypic block hypertensive diseases (I10-I15) and 17 SNPs are associated with both hypertensive diseases (I10-I15) and Ischaemic heart diseases (I20-I25). **S6 Fig** shows the associations between SNPs (red circle) and the set of malignant neoplasms phenotypic blocks (C00-C97; blue square) identified by the HCLC-FC method. There are a total of 15 phenotypic blocks within the set of the malignant neoplasms (C00-C97), such as melanoma and other malignant neoplasms of skin (C43-C44) and malignant neoplasms of mesothelial and soft tissue (C45-C49).

**Fig 3.**
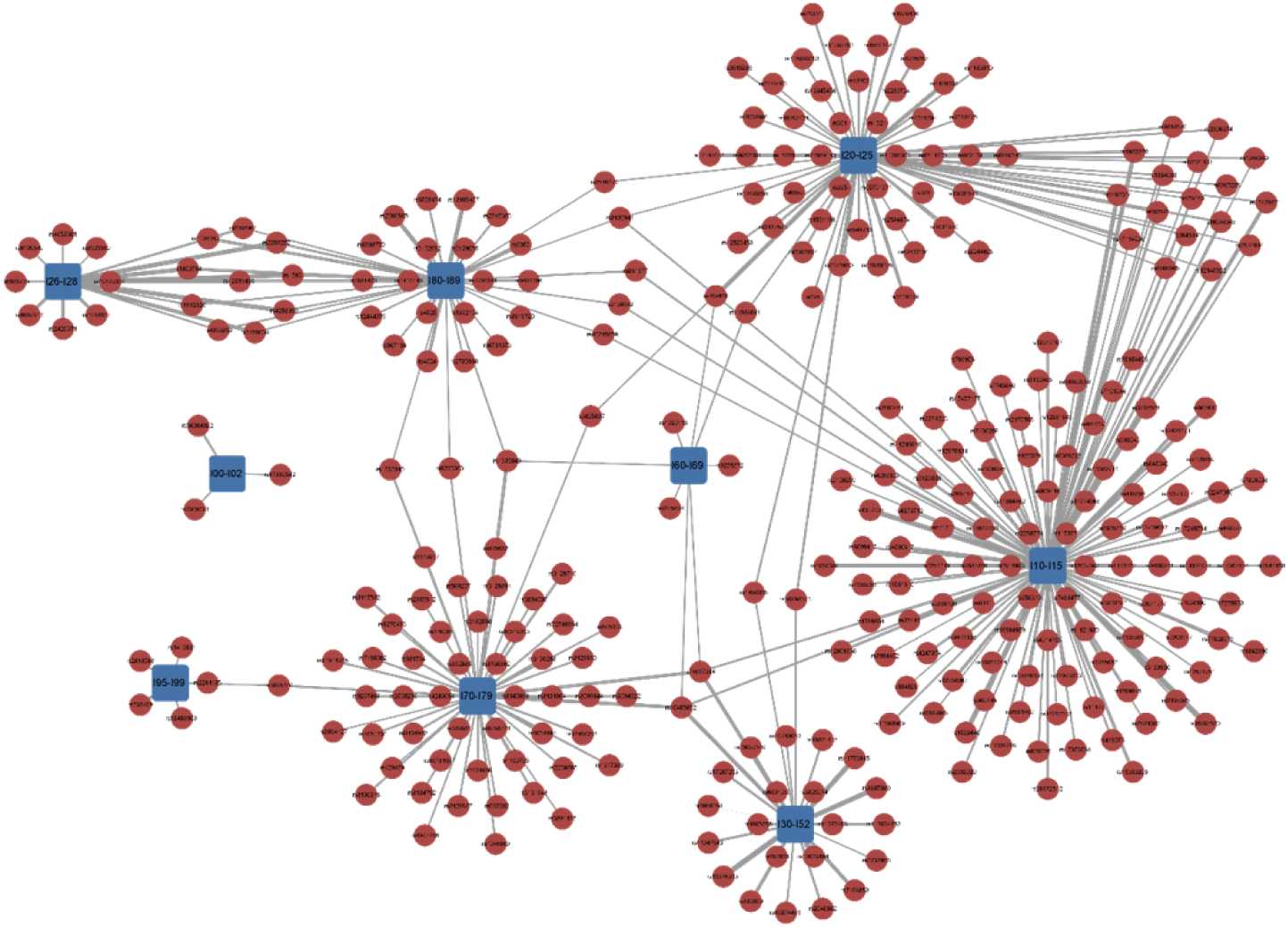
The associations between SNPs and the circulatory system phenotypic blocks identified by the HCLC-FC method. The red circles represent SNPs, and the blue squares represent the set of diseases of the circulatory system phenotypic blocks I00-I99. The width of the connection line represents the strength of association (-log10 scale p-value).

## Discussion

GWAS have become a very effective research tool to investigate associations between genetic variation and a disease/phenotype. In spite of the success of GWAS in identifying thousands of reproducible associations between genetic variants and complex diseases, in general, the association between genetic variants and a single phenotype is usually weak. It is increasingly recognized that joint analysis of multiple phenotypes can be potentially more powerful than the univariate analysis and can shed new light on underlying biological mechanisms of complex diseases. As a complementary approach to GWAS, PheWAS analyze many phenotypes with a genetic variant and combine both the exploration of phenotypic structure and genotypic variation [13]. Similar to the widely used GWAS approaches, existing methods for PheWAS largely focus on the association between a single genetic variant with a large number of candidate phenotypes and test the association between one genetic variant and one phenotype at a time.

In this paper, we develop a novel and powerful multivariate method, HCLC-FC, to test the association between a genetic variant with multiple phenotypes in each phenotypic category. HCLC-FC involves three steps. In the first step, we use the bottom-up hierarchical clustering method [24] to partition a large number of phenotypes into disjoint clusters within each category. In the second step, we apply the clustering linear combination method [22] to combine test statistics within each category based on the phenotypic clusters and obtain a p-value from each phenotypic category. In the third step, we propose a large-scale association testing procedure with theoretical guarantees for FDR control under flexible correlation structures. We perform extensive simulation studies to compare the performance of HCLC-FC with that of other existing methods. The results show that the existing methods using our proposed FDR control procedure can control FDR at a nominal level, and our proposed HCLC-FC method outperforms the other three methods we compare with under the six models for different within-group and between-group phenotypic correlation structures. Finally, we also evaluate the performance of HCLC-FC through a set of 1869 case-control phenotypes based on ICD-10 code across more than 300,000 samples from the UK Biobank, where these phenotypes can be classified into 260 ICD-10 level 1 blocks. The real data analysis results show that HCLC-FC not only can well-control type I error rates but also can identify 1,244 SNPs that have previously been reported to be associated with some phenotypes in the GWAS catalog. In this analysis, we classify phenotypes according to the ICD-10 code. Future analyses using genetic architecture or disease network that incorporate genotypic and phenotypic information to classify the whole phenome into numerous phenotypic categories [17, 19, 20] may improve the performance of the proposed method.

In summary, HCLC-FC has several important advantages over other existing methods for association studies using multiple phenotypes. First, it clusters phenotypes within each phenotypic category, which reduces the degrees of freedom of the association tests and has the potential to increase statistical power. Second, it is computationally fast and easy to implement. The CLC approach [22] uses a simulation procedure to estimate the p-value of the final test statistic. HCLC-FC has an asymptotic distribution which avoids the computational burden of simulation. Third, the newly developed FDR controlling process is based on p-values and does not depend on test statistics. Therefore, it is more general and can be applied to other multiple testing procedures to control FDR.

## Acknowledgements

Part of this research has been conducted using the UK Biobank Resource under application number 41722 and the NHGRI-EBI GWAS Catalog. Superior, a high-performance computing infrastructure at Michigan Technological University, was used in obtaining results presented in this publication.

## Author contributions

S. Z. and Q. S. designed research, X. L. and S. Z. performed the statistical analysis, X. C. performed real data analysis, and X. L., X. C., Q. S., and S. Z. wrote the manuscript.

## Supplementary Materials

**Fig S1.**
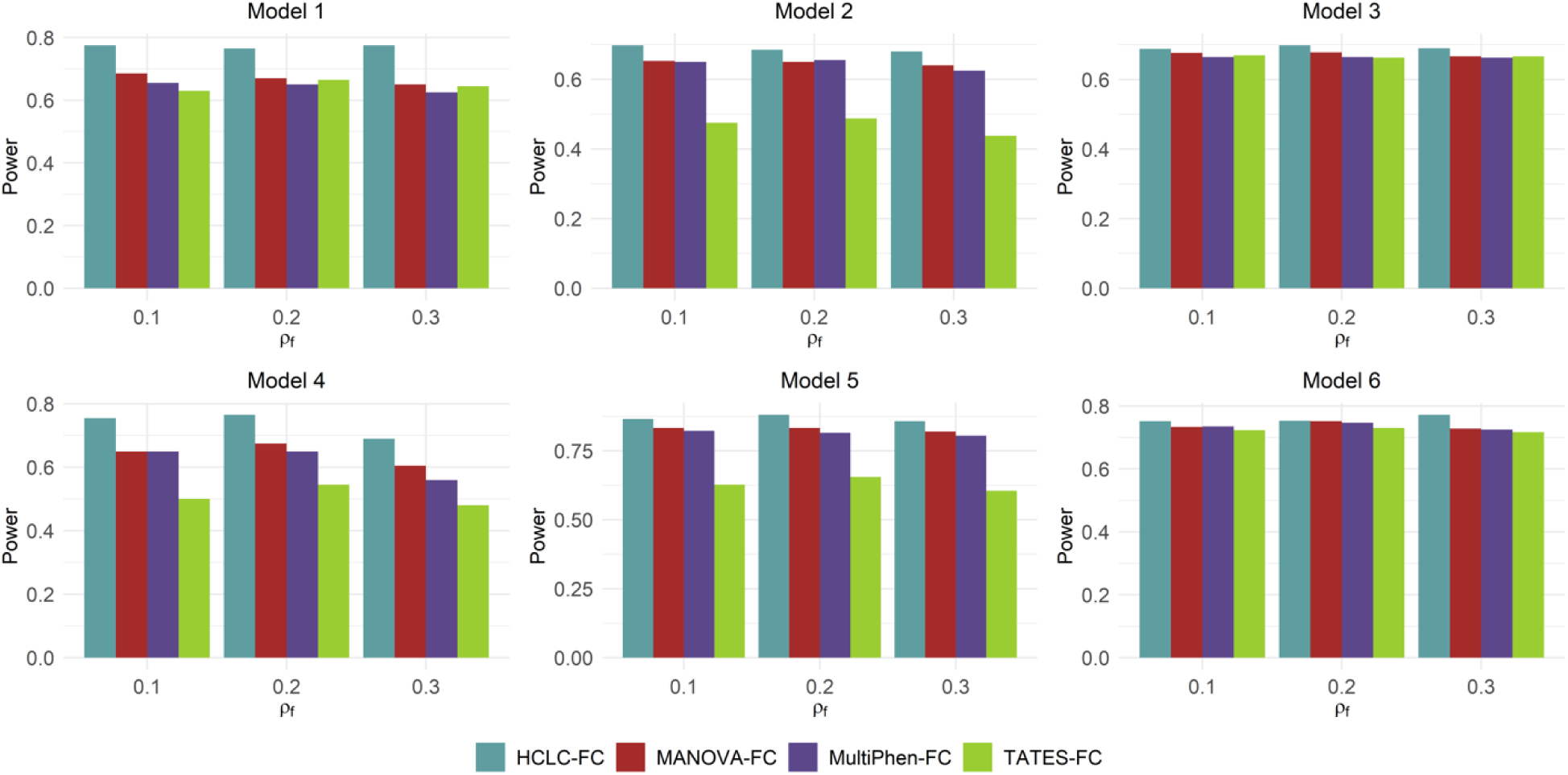
Power comparisons of the four tests for the power as a function of correlation between phenotypic categories (*ρ*_*f*_) under the six models for 1,000 phenotypes (*K* = 1,000). MAF is 0.3. The sample size (*n*) is 2,000. *ρ*_*e*_ = 0.3 and *c*^2^ = 0.5. The effect sizes of the six models are 0.014, 0.060, 0.090, 0.006, 0.060, and 0.090. The power of the four tests is evaluated using 200 replicated samples at a nominal FDR level of 5%.

**Fig S2.**
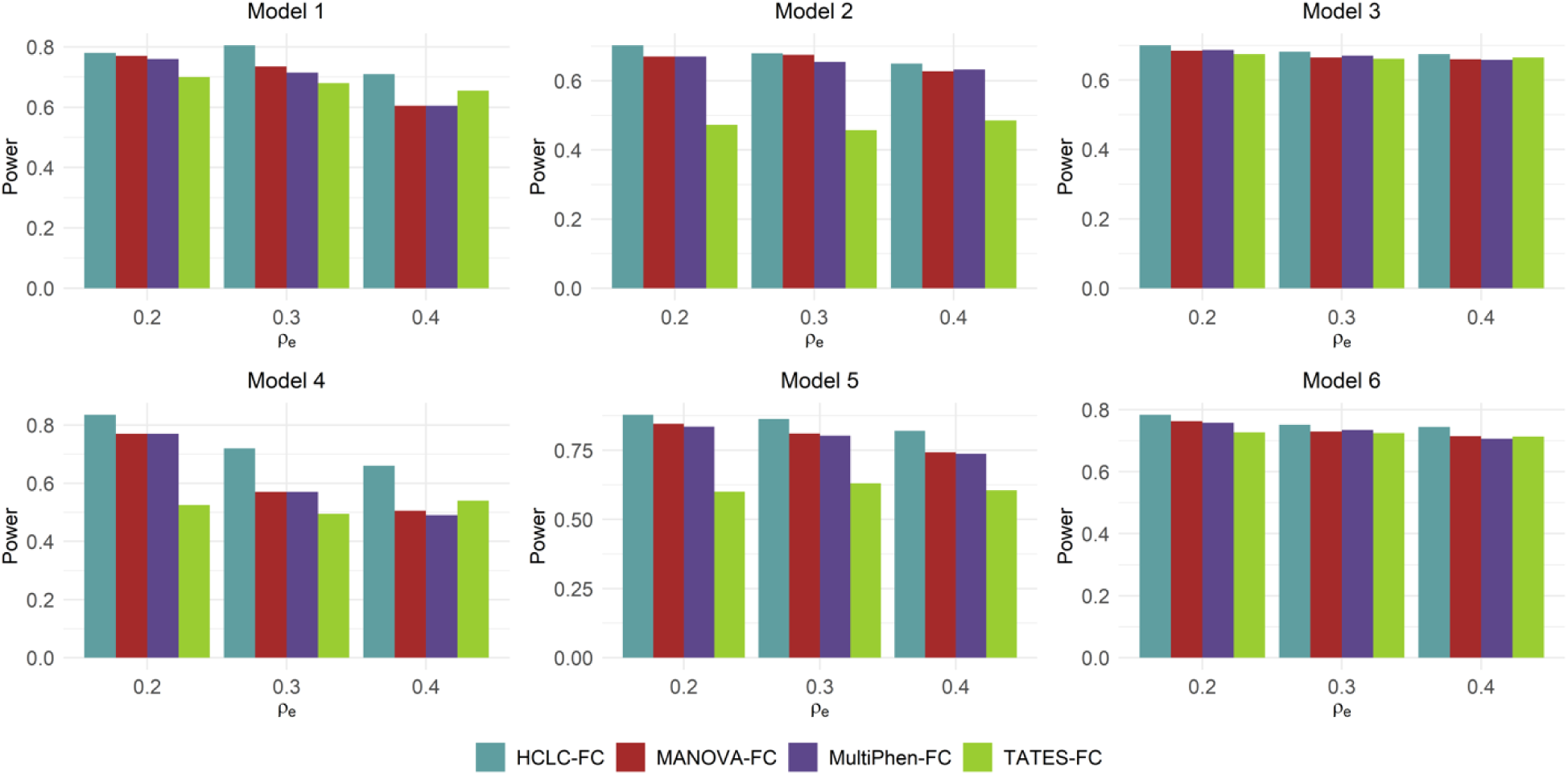
Power comparisons of the four tests for the power as a function of correlation within each phenotypic category (*ρ*_*e*_) under the six models for 1,000 phenotypes (*K* = 1,000). MAF is 0.3. The sample size (*n*) is 2,000. *ρ*_*f*_ = 0.2 and *c*^2^ = 0.5. The effect sizes of the six models are 0.014, 0.060, 0.090, 0.006, 0.060, and 0.090. The power of the four tests is evaluated using 200 replicated samples at a nominal FDR level of 5%.

**Fig S3.**
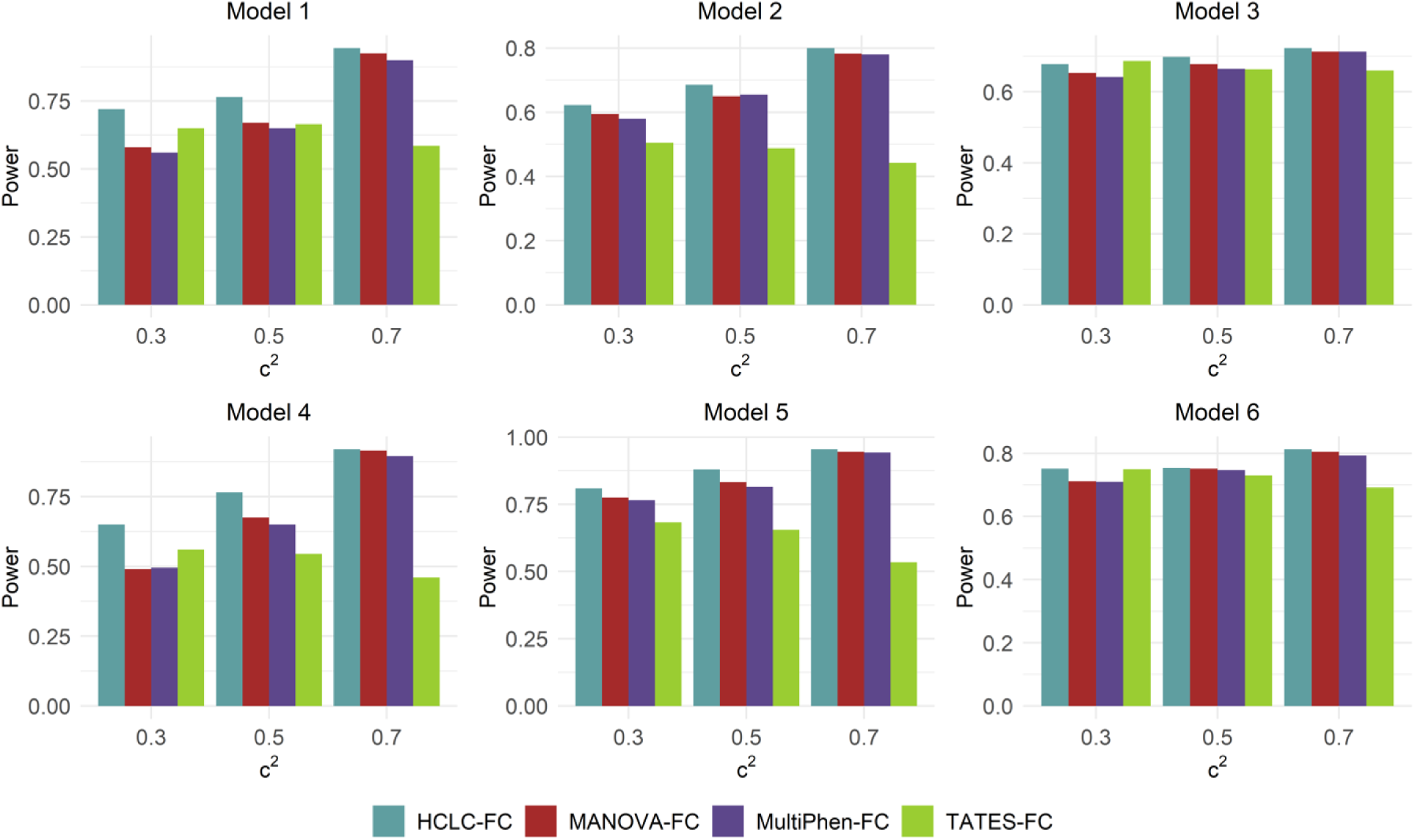
Power comparisons of the four tests for the power as a function of (*c*^2^) under the six models for 1,000 phenotypes (*K* = 1,000). MAF is 0.3. The sample size (*n*) is 2,000. *ρ*_*f*_ = 0.2 and *ρ*_*e*_ = 0.3. The effect sizes of the six models are 0.014, 0.060, 0.090, 0.006, 0.060, and 0.090. The power of the four tests is evaluated using 200 replicated samples at a nominal FDR level of 5%.

**Fig S4.**
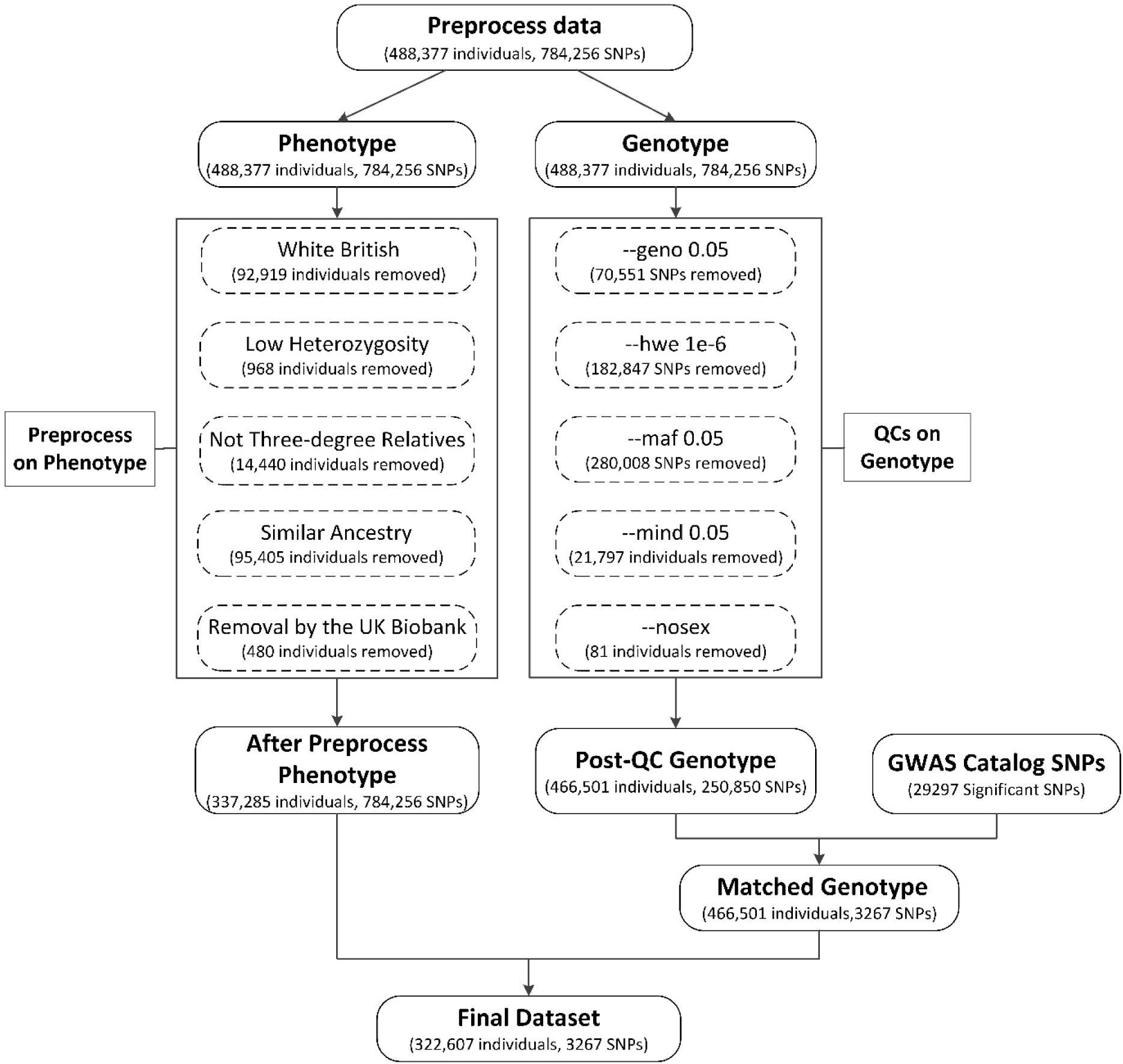
Flow chart of UK Biobank data preprocessing. *Preprocess on phenotype*: i. Select White British subjects (White British); ii. Remove individuals who are marked as outliers for heterozygosity or missing rates (Low Heterozygosity); iii. Exclude individuals who have been identified to have ten or more third-degree relatives or closer (Not Three-degree Relatives); iv. Remove individuals having very similar ancestry based on the principal component analysis of the genotypes (Similar Ancestry); v. Remove individuals that are recommended for removal by the UK Biobank (Removal by the UK Biobank). *Quality controls (QCs) on genotype*: Filter out genetic variants, with i. Missing rate larger than 5% (“--mind 0.05”), ii. Hardy-Weinberg equilibrium exact test p-values less than 10^−6^ (“--hwe 1e-6”), iii. Minor allele frequency (MAF) less than 5% (“--maf 0.05”). We also filter out individuals, with iv. Missing rate larger than 5% (“--mind 0.05”) v. Individuals without sex (“--no-sex”).

**Fig S5.**
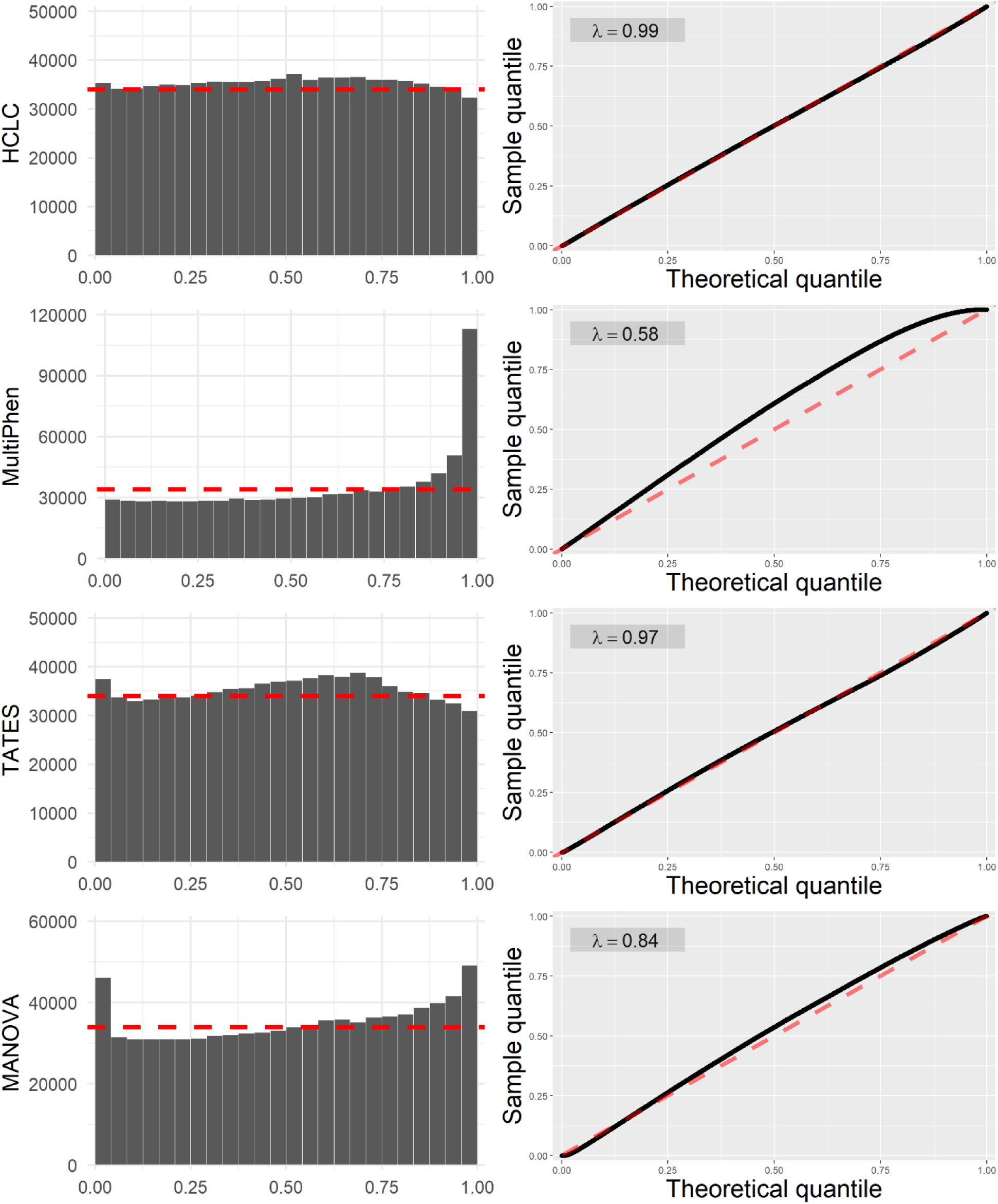
The histogram of p-values (left) and QQ plot for uniform distribution of each method (right) based on 3,267 replicates. The red dashed line represents the theoretical frequency and quantile for the standard uniform distribution.

**Fig S6.**
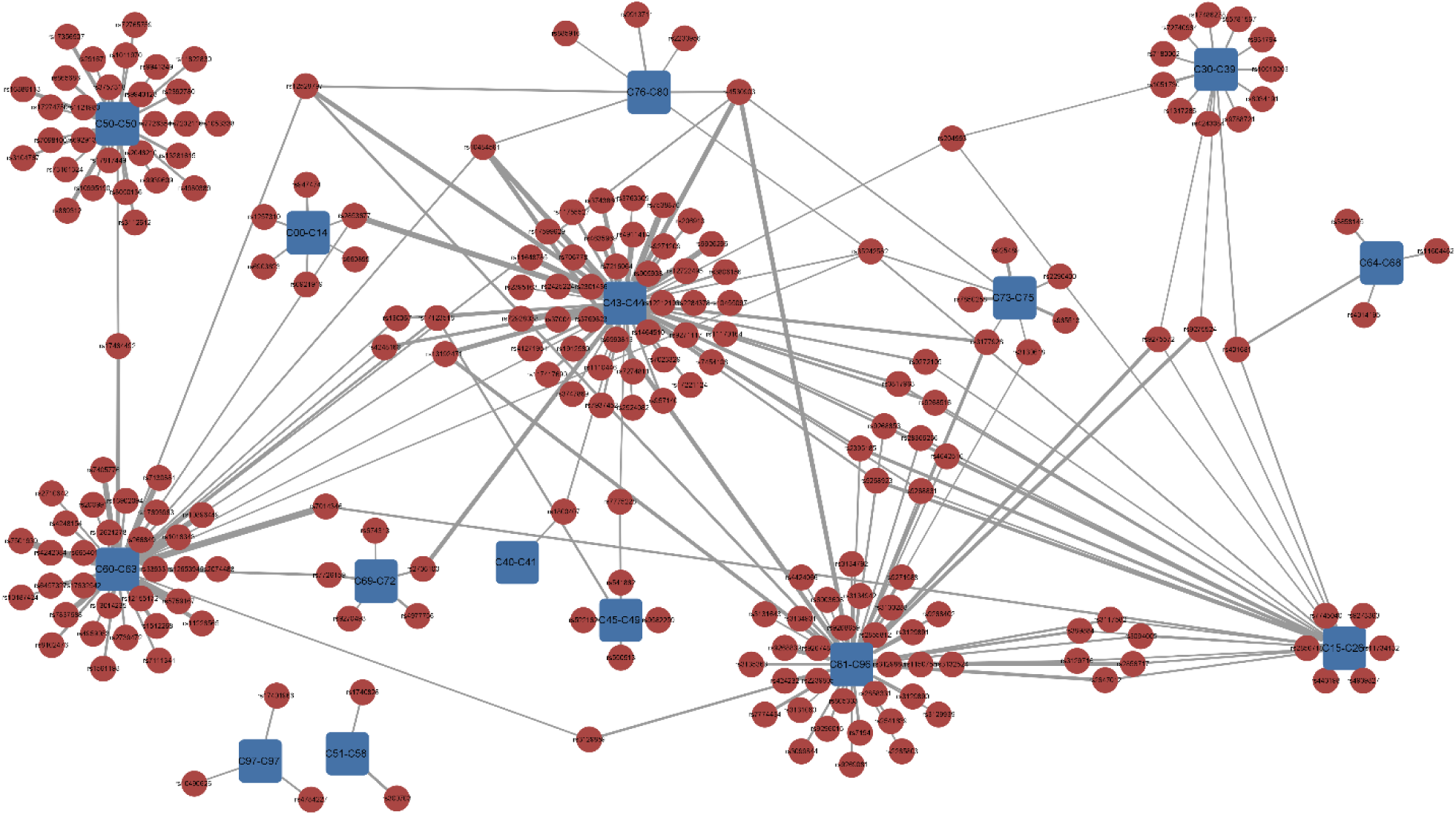
The associations between SNPs and the malignant neoplasms phenotypic blocks identified by the HCLC-FC method. The red circles represent SNPs, and the blue squares represent the set of malignant neoplasms phenotypic blocks C00-C97. The width of the connection line represents the strength of association (-log10 scale p-value).

